# Characterisation of fasting and postprandial NMR metabolites: insights from the ZOE PREDICT 1 Study

**DOI:** 10.1101/2022.11.14.516406

**Authors:** Kate M Bermingham, Mohsen Mazidi, Paul W Franks, Tyler Maher, Ana M Valdes, Inbar Linenberg, Jonathan Wolf, George Hadjigeorgiou, Tim D Spector, Cristina Menni, Jose M Ordovas, Sarah E Berry, Wendy L Hall

**Author notes:** Equal contribution (Kate M Bermingham, Mohsen Mazidi, Sarah E Berry, Wendy L Hall). Corresponding author, **Corresponding author information:** Sarah E Berry, PhD Department of Nutritional Sciences, King’s College London, London, United Kingdom.

## Abstract

**Background:** Postprandial metabolomic profiles and their inter-individual variability are not well characterised. Here we describe postprandial metabolite changes, their correlations with fasting values and their inter- and intra-individual variability following a standardised meal in the ZOE PREDICT 1 cohort.

**Methods:** In the ZOE PREDICT 1 study (*n* = 1,002 (NCT03479866)), 250 metabolites, mainly lipids, were measured by Nightingale NMR panel in fasting and postprandial (4 and 6 h after a 3.7 MJ mixed nutrient meal, with a second 2.2 MJ mixed nutrient meal at 4 h) serum samples. For each metabolite, inter- and intra-individual variability over-time was evaluated using linear mixed modelling and intraclass-correlation coefficients (ICC) calculated.

**Results:** Postprandially, 85% (of 250 metabolites) significantly changed from fasting at 6h (47% increased, 53% decreased; Kruskal-Wallis), with 37 measures increasing by >25%, and 14 increasing by >50%. The largest changes were observed in very large lipoprotein particles and ketone bodies. Seventy-one percent of circulating metabolites were strongly correlated (Spearman’s rho >0.80) between fasting and postprandial timepoints, and 5% were weakly correlated (rho <0.50). The median ICC of the 250 metabolites was 0.91 (range 0.08-0.99). The lowest ICCs (ICC<0.40, 4% of measures) were found for glucose, pyruvate, ketone bodies (β-hydroxybutyrate, acetoacetate, acetate) and lactate.

**Conclusions:** In this large-scale postprandial metabolomic study, circulating metabolites were highly variable between individuals following a mixed challenge meal. Findings suggest that a meal challenge may yield postprandial responses divergent from fasting measures, specifically for glycolysis, essential amino acid, ketone body and lipoprotein size metabolites.

## Introduction

Advancements in metabolomics and the development of comprehensive high-throughput profiling have enabled the simultaneous quantification of multiple biomarkers in large cohorts (1–4). This progressed our understanding of mechanistic pathways linking metabolites to disease risk and enabled early identification of elevated risk for early atherosclerosis, type 2 diabetes, diabetic nephropathy, cardiovascular diseases, and all-cause mortality (5). To date, metabolomic profiles have been reported mainly in the fasting state (1). However, the physiological relevance of fasting analyses is a point of debate (6–7), since we consume multiple mixed-nutrient meals throughout the day, and therefore spend most of our time in the highly dynamic postprandial state.

Moreover, postprandial metabolic dysregulation is an independent risk factor for non-communicable diseases (8–10), but the relevance for health of non-standard meal-induced postprandial metabolomic markers is less clear. Standard clinical biochemistry analysis of blood glucose, TG and insulin alone does not fully represent the multiple downstream postprandial metabolic changes that can be captured from metabolomic analysis and potentially harnessed for improved sensitivity in the prediction of pre-clinical risk of cardiometabolic diseases. To date, studies examining postprandial metabolomics have been conducted in small cohorts (11), or focused on specific metabolites instead of quantifying a broad range of metabolomic responses (12, 13). Furthermore, despite growing awareness of the large inter-individual variability in metabolic responses to food (14), this has rarely been explored beyond simple clinical measures.

The ZOE PREDICT 1 study was designed to quantify and predict individual variations in fasting and postprandial TG, glucose and insulin responses to sequential standardised meals in a tightly controlled setting (14). The aim of this study was to explore and compare inter-individual fasting and postprandial variabilities in metabolomic profiles.

## Materials and Methods

The ZOE PREDICT 1 study (NCT03479866) was a single-arm, single-blinded study (June 2018 to May 2019) in 1,102 healthy adults, aged 18-65 y (*n*=1,002 from the United Kingdom (UK); for the full protocol, see Berry *et al*. (15). The study consisted of a 1-day clinical visit at baseline followed by a 13-day at-home period, although this paper only focuses on the 1-day clinical visit. Primary outcomes are reported elsewhere (13, 14, 16). Secondary outcome metabolomic data measured by NMR (at the baseline visit only) is reported in this paper. At baseline (day 0), participants arrived fasted and were given a standardised metabolic challenge meal for breakfast (0h; 86g carbohydrate, 53g fat, 16g protein; 3.7MJ) and a test lunch (4h; 71g carbohydrate, 22g fat, 10g protein; 2.2MJ). The fat was high oleic sunflower oil; 85% oleic acid (18:1n-9) and 8% linoleic acid (18:2n-6). Fasting and postprandial (0-6h) venous blood was collected to determine concentrations of serum glucose, insulin, TG and metabolomics (using NMR described below). The trial was approved in the UK by the Research Ethics Committee and Integrated Research Application System (IRAS 236407), registered on ClinicalTrials.gov (NCT03479866) and was run in accordance with the Declaration of Helsinki and Good Clinical Practice.

### Metabolite measurements

Metabolite concentrations were quantified at three time points from serum at fasting, 4h and 6h postprandially using high-throughput NMR metabolomics (2020 Platform; Nightingale Health, Helsinki, Finland). The metabolomics platform provides 250 parameters (concentrations, ratios, size, percentages) derived from 163 raw metabolite measures (concentrations and size). Details of the experimentation and epidemiological applications of the NMR metabolomics platform have been reviewed previously (17).

### Statistical analysis

Statistical analysis was performed in the R environment for statistical computing version 3.5.1 (R Foundation for Statistical Computing, Vienna, Austria. https://www.R-project.org/). Metabolites were characterised by mean, median, 25th and 75th percentiles at fasting and 4h and 6h postprandially. Absolute change and percentage change were calculated. Kruskal-Wallis tests were performed to evaluate differences in the median concentrations at fasting and 6h. Spearman’s correlation assessed the relationship between measures at fasting and 6h and the Fligner-Killeen test compared variances at fasting and 6h. Time-dependent changes in metabolite concentrations within individuals were evaluated using mixed models. Total variance in plasma metabolites was decomposed into interindividual variance, which can also be considered the variance of the usual level in a population and intraindividual variance, which reflects variability around the usual level within an individual. Fasting and postprandial metabolite levels were included as the outcome variables, with time as a fixed effect and participant ID as a random effect. Intraclass correlations (ICC) were calculated, denoting the proportion of the population’s biologic variability that is due to the interindividual variation (18, 19). A high ICC can be obtained by low intra- and/or high inter-individual variance. A low ICC is attributable to high intra- and/or low inter-individual variance. The Benjamini-Hochberg correction for multiple comparisons was applied (20). Statistically significant thresholds were based on FDR cut-offs (*q*< 0.05).

## Results

1,002 generally healthy adults completed baseline (day 0) measurements and the sequential test meal challenge. Descriptive characteristics of study participants are summarized in [Supplementary Table 1](https://docs.google.com/spreadsheets/d/1mBWYdNx2eCas_waUv9Eg2q5wAhyDCQ-9uCv41tE080c/edit#gid=1133978210) and the study design is shown in **Figure 1**. Participants were aged between 18.5 and 65.9 (mean 45.6 ± 11.9) years with a mean BMI of 25.6 (± 5.0) kg/m^2^.

**Figure 1.**
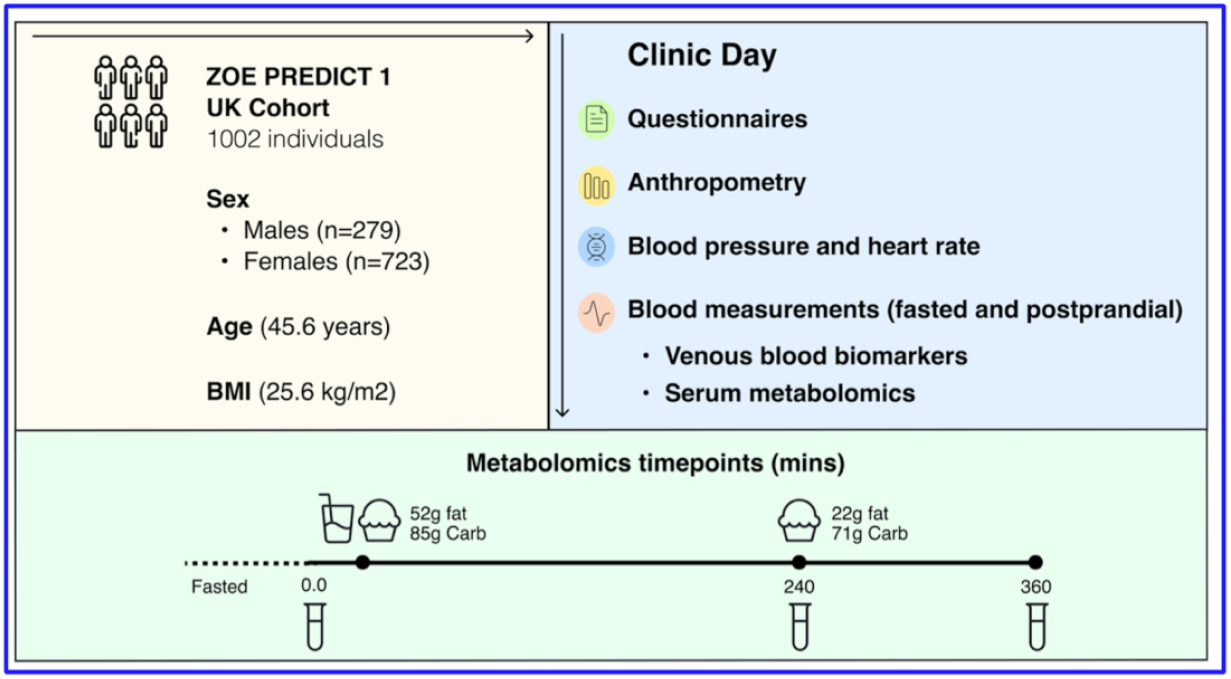
ZOE PREDICT 1 Study Design. Participants arrived fasted for their baseline visit and were given a standardised breakfast (0h, metabolic challenge meal, 86g carbohydrate, 53g fat) and lunch (4h, 71g carbohydrate, 22g fat). Concentrations of glucose, TG and NMR metabolites were determined from venous blood collected at multiple timepoints postprandially. Anthropometric and fasting biochemistry measurements were also measured.

### Characterization of metabolite biomarkers

Metabolite concentrations measured as mean, median and IQR are reported in [Supplementary Table 2](https://docs.google.com/spreadsheets/d/1mBWYdNx2eCas_waUv9Eg2q5wAhyDCQ-9uCv41tE080c/edit#gid=1133978210) for fasting, 4 and 6h values for 250 metabolites. A selection of metabolites (*n*=75) is also presented in **Table 1**

**Table 1.**
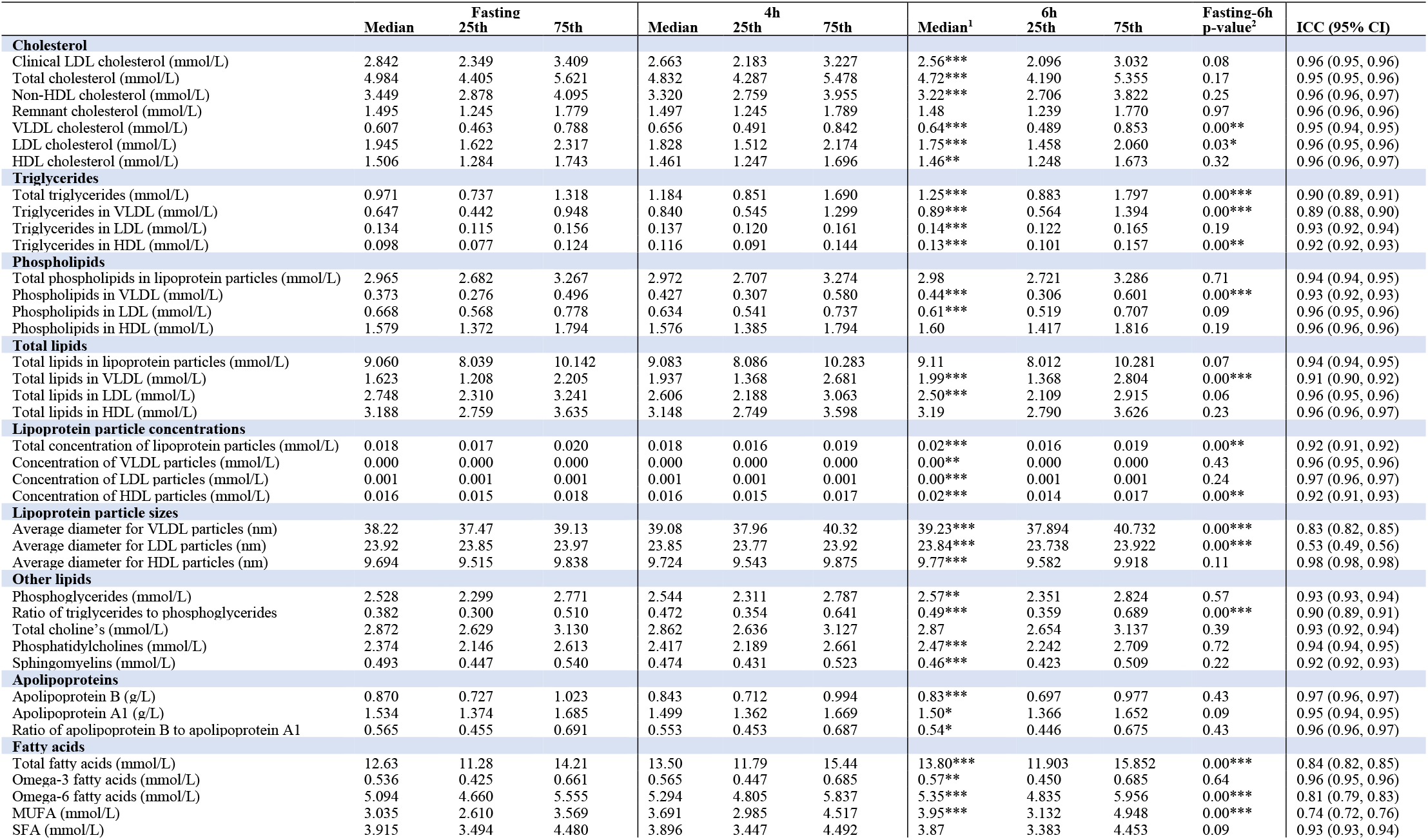
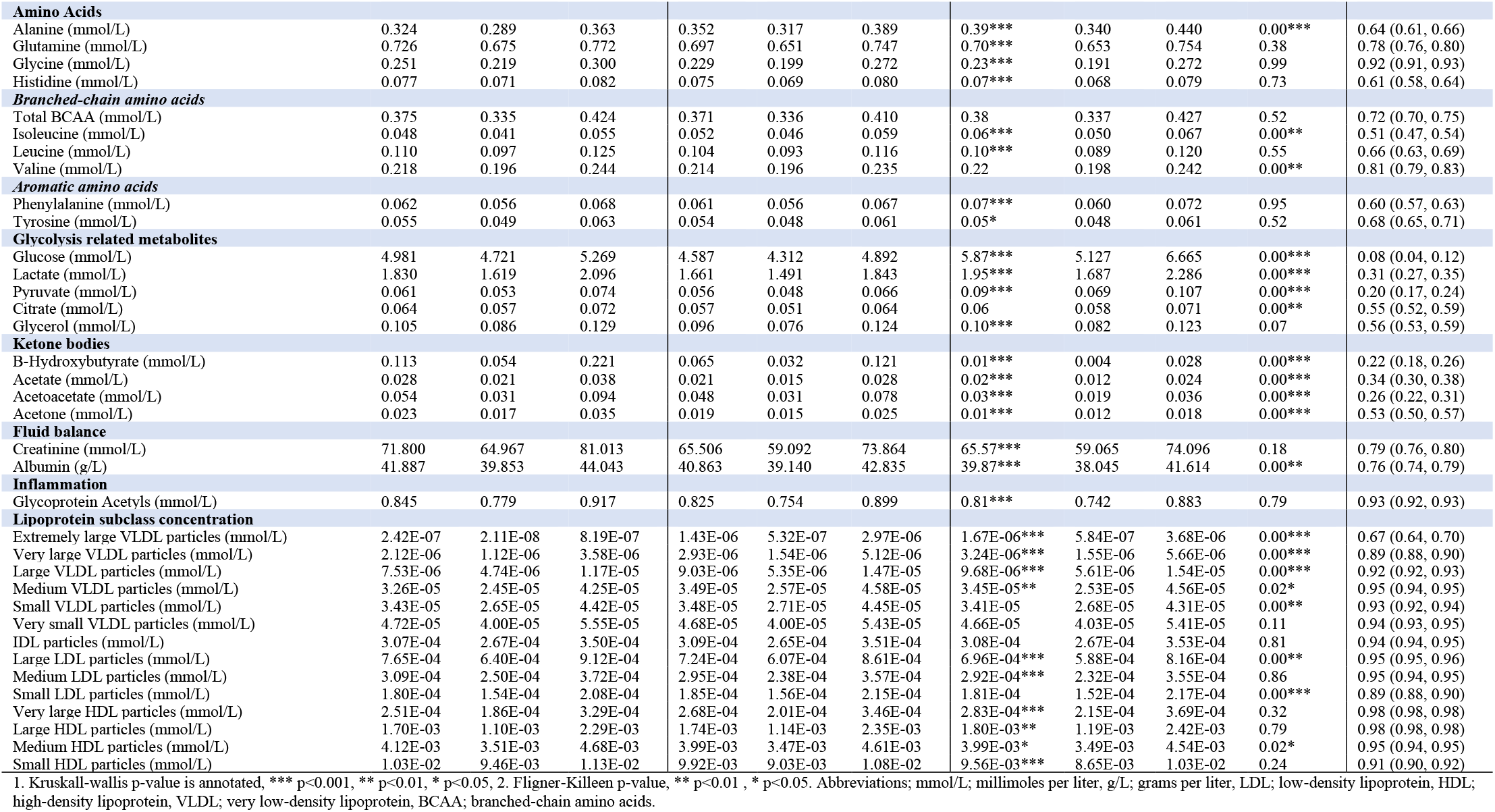
Characterization of concentrations of the metabolomic markers

### Postprandial change

Postprandially, 85% of the 250 total metabolite outcomes measured had a significant absolute change at 6h from fasting (47% with a significant increase and 53% with a significant decrease; Kruskal-Wallis FDR<0.05). At 6h, 37 of the 250 metabolites changed by >25% from fasting values (30 increased and 7 decreased by >25%), of which 14 changed by >50% from fasting values (12 increased and 2 decreased by >50%) (**Supplementary Table 2**; median % change). The largest postprandial increases (median % change; 0-6h) were elicited in the XXL-VLDL particles, specifically particle number (XXL-VLDL-P; 440%), TG (XXL-VLDL-TG; 676%), phospholipid (XXL-VLDL-PL; 570%) and total lipid (XXL-VLDL-L; 379%) concentrations. The largest postprandial decreases (median % change; 0-6h) were observed in ketone bodies (β-hydroxybutyrate: −85%, acetoacetate: −49%, acetate: −40%, acetone: −400%) and the percentage contribution of cholesterol (esters (CE) and total (C)) to XL-VLDL (XL-VLDL-CE: −57%, XL-VLDL-C: **-**46%). Traditional clinical measures (TG, glucose and non-HDL), lipoprotein particle sizes (due to their strong association with disease risk) and the variables with the largest postprandial change (>25%; 0-6h), within each class of metabolite, are shown in **Figure 2** and **Figure 3**.

**Figure 2.**
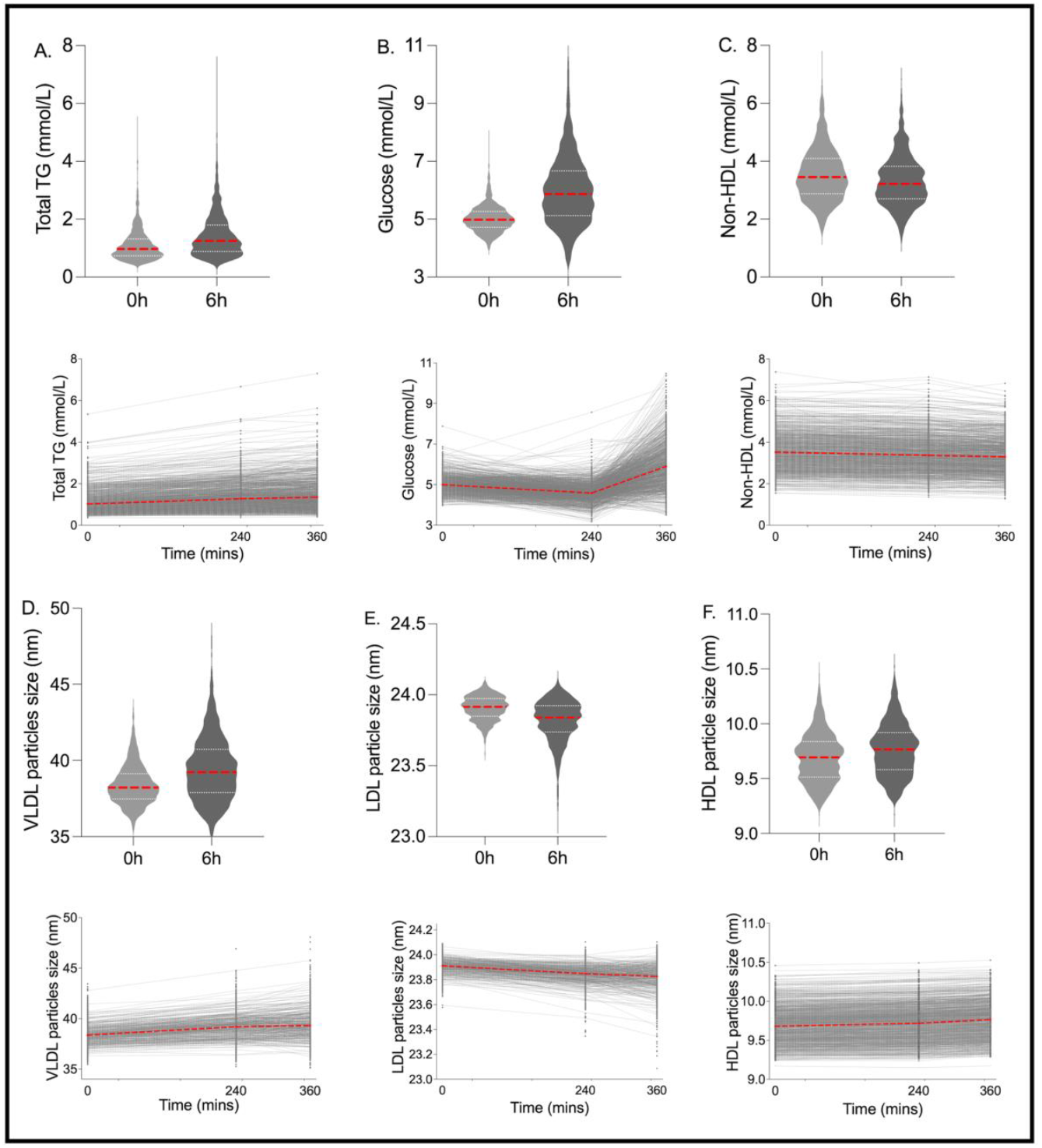
Inter-individual variation and distribution for traditional clinical metabolites and lipoprotein particle size. Fasting and postprandial concentrations of **A.** triglycerides (mmol/L), **B.** glucose (mmol/L), **C.** non-HDL (mmol/L); and particle sizes of: **D. V**LDL (nm), **E.** LDL (nm), **F.** HDL (nm). n=1002.

**Figure 3.**
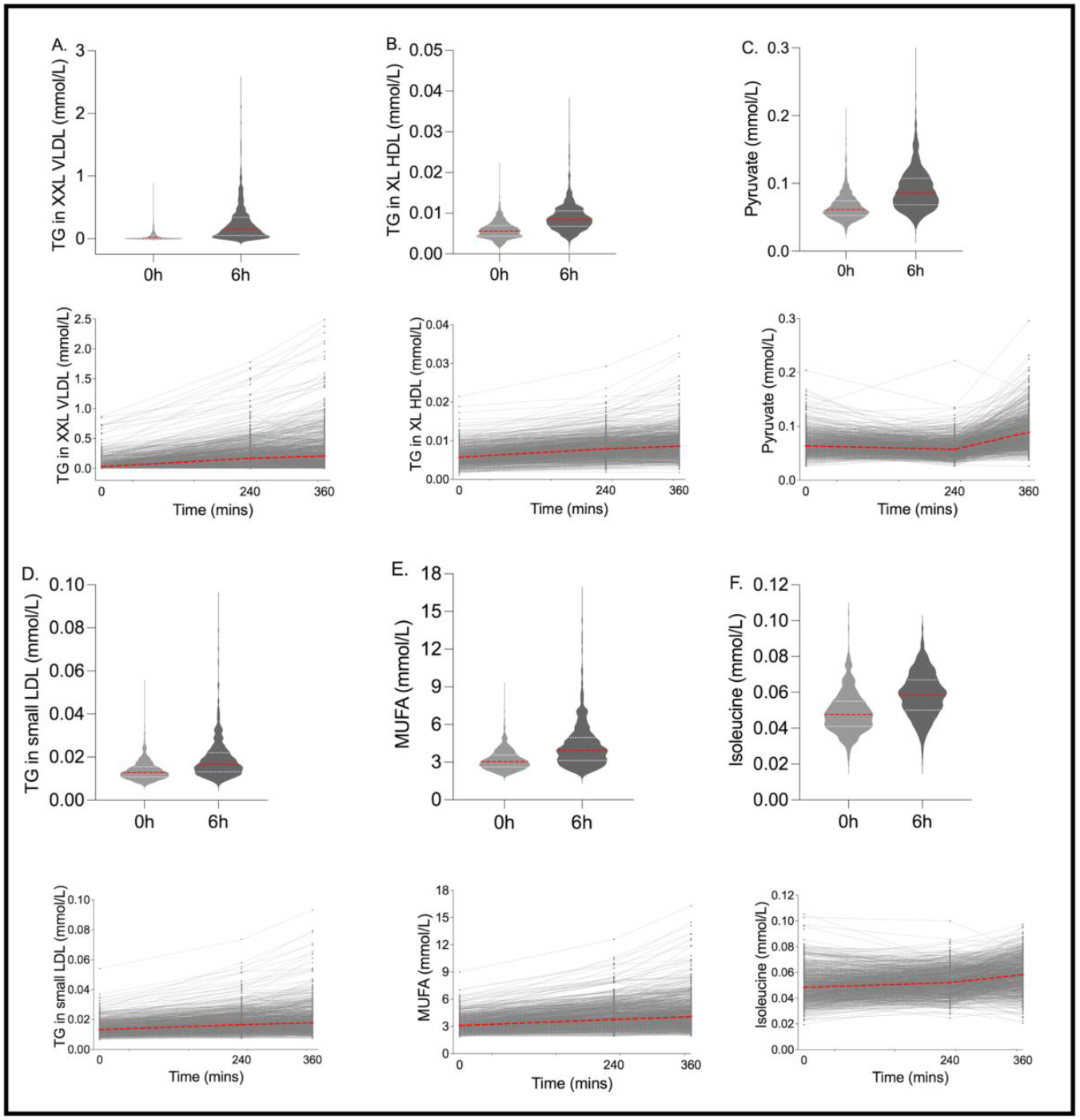
Metabolites with the greatest postprandial change and/or postprandial inter-individual variability. Fasting and postprandial concentrations of **A.** triglycerides in extremely large VLDL particles and chylomicrons (TG in XXL VLDL), **B.** triglycerides in large HDL particles and chylomicrons (TG in XL HDL), **C.** triglycerides in LDL particles and chylomicrons (TG in LDL), **D.** Pyruvate, **E.** Mono-unsaturated fatty acids (MUFA), **F.** Isoleucine. n=1002.

### Correlation between fasting and postprandial metabolites

Postprandial concentrations of key food-induced metabolic markers, glucose and TG, are known to be more discriminatory of CVD risk than their fasting values (8–10). However, if postprandial metabolites are closely correlated to their fasting values, there is minimal utility in conducting burdensome postprandial studies. Therefore, we assessed the correlation between fasting and postprandial measures to explore the value of measuring non-standard clinical measures postprandially. For most measures, the 6h values were strongly correlated with fasting values (Spearman’s rank correlation coefficient >0.80 in 71% of measures; **Supplementary Table 2**); however, low correlations (rho<0.50) were observed for 5% of measures including, ketone bodies (β-hydroxybutyrate, acetate, acetoacetate), as well as glucose, pyruvate, lactate, LDL diameter, isoleucine and phenylalanine. The lack of correlation for these measures may be due to significant variation within individuals (differences from one time point to another).

### Interindividual variability in metabolites over time

Given the highly variable postprandial responses observed in traditional clinical measures (TG, glucose and insulin) following a standardized meal in healthy individuals (15), we explored the variability in postprandial metabolomic responses. The proportion of total variability attributable to between-subject differences, as determined by the ICC (ratio of between-person variance and total variance (sum of between- and within-person variances)), was high for most metabolites (ICC?0.75; 83%, 0.51-0.74; 12%, 0.40-0.50; 1%, <0.40; 4%). The median ICC of the 250 metabolites was 0.91 (range 0.08-0.99). The metabolites with the highest ranked ICCs included HDL measures (cholesterol, cholesterol esters, particle concentration, total lipids, phospholipids and free-cholesterol in very large and large HDL), apolipoprotein B, the ratio of apolipoprotein B to apolipoprotein A, and concentration of LDL particles. A selection of metabolites had lower ICCs (<0.40), meaning variation around an individual’s usual level was larger. These metabolites include glucose, pyruvate, ketone bodies (β-hydroxybutyrate, acetoacetate, acetate) and lactate.

The inter-individual pattern of response for fasting and 6h time-points was also assessed using the Fligner-Killeen test of variance. There were large differences in the variance of the data at 6h versus fasting (Fligner-Killeen test of variance p<0.001 for 39% of measures; p<0.01 for 49% of measures; p<0.05 for 58% of measures, **Supplementary Table 2**), illustrating the differential variability (spread) in the postprandial versus fasting state (**Figure 2, 3**).

## Discussion

This study set out to describe meal-induced changes in metabolomic markers, and to compare fasting and postprandial correlations and differences in inter-individual variability. Most non-traditional clinical metabolites from the Nightingale NMR panel showed large inter-individual variability following a mixed challenge meal. Greater inter-individual variability was observed in traditional clinical postprandial measures relative to equivalent fasting measures. A lack of correlation over time between fasting and postprandial metabolite concentrations was due to significant variation within individuals (differences from one time point to another). These findings suggest that postprandial responses for glycolysis, essential amino acid, ketone body and lipoprotein size metabolites may provide more insight into disease risk than fasting measures alone.

The metabolomic composition of plasma is affected by many factors including diet, sex, age, genetics, lifestyle, and the gut microbiome (18, 21–23). Research demonstrates that certain metabolites vary within an individual; for example plasma 1H NMR metabolites vary during the menstrual cycle within pre-menopausal females (24). Others are more stable over longer periods of time, reflecting usual levels necessary for large-scale epidemiological research. The reliability over time of fasting blood metabolites has been investigated (18, 19, 25) and reliability over time has been shown to decrease in non-fasting samples (22). Food intake influences the metabolomic profile, but short-term postprandial metabolomic responses, specifically in lipids and their subclasses, are less understood. This research shows that a meal challenge yielded lower ICCs in glycolysis, essential amino acid, ketone body and lipoprotein size metabolites. These measures have higher intra-individual variability and thus may provide more insight into divergent metabolic responses and associated disease risk. Most fasting and postprandial metabolites measured by the Nightingale NMR panel, mainly lipids and their subclasses, were shown to be stable in the postprandial response phase. Thus, analysis by more comprehensive metabolic panels may reveal postprandial perturbations not detected in this panel.

The postprandial percentage changes were greatest in the VLDL parameters, particularly in the concentration and lipids of the largest VLDL particles, in agreement with previous studies (26) and likely a marker of exogenous chylomicron TG. TG-rich lipoproteins, chylomicrons and VLDL, and their remnants (all captured in this platform under the VLDL particles), increase in the circulation following a fatty meal and are known to be atherogenic, with non-fasting TG concentrations being strongly associated with risk of CHD, stroke, and mortality (27), and non-fasting small and large VLDL-C accounting for 40% of increased risk of myocardial infarction associated with higher BMI (28). Elevated postprandial TG concentrations that persist for up to 6 hours and beyond are mainly attributable to greater increases in subclasses of large VLDL (29). Our previous work has shown that using postprandial plasma TG concentrations as an indicator of the atherogenic potential of different meals may be misleading since a saturated fat-rich meal-induced lower postprandial TG concentrations, but higher large VLDL concentrations at 6-8h, compared with a monounsaturated-rich oil (30). Therefore, quantifying postprandial large VLDL particles and particle composition may be more discriminatory than total TGs when assessing the atherogenic potential of a meal or food and their implications for CVD risk.

The strengths of this study are the large study population, repeated postprandial timepoints allowing analysis at peak lipemia and the later postprandial phase, and the design of the postprandial challenge, which adopted physiologically-relevant macronutrient profiles and sequential meals. However, limitations of the study included a lack of longer postprandial follow-up (up to 8-10 h) which may have revealed NMR measures that were more discriminatory of metabolic status and the limited panel of metabolites measured. We also could not partition technical and intra-individual variability across timepoints.

In conclusion, this paper provides a large, comprehensive NMR spectroscopy metabolomics resource for lipid and postprandial metabolic research and demonstrates that postprandial responses for glycolysis, essential amino acid, ketone body and lipoprotein size plasma metabolites may provide more insight into favourable metabolic responses and associated disease risk than fasting measures alone.

## Abbreviations

BCAA: branched-chain amino acids
C: cholesterol
CE: cholesterol ester
FC: free cholesterol
GlycA: glycoprotein acetyls
HDL: high-density lipoprotein
HOMA-IR: homeostatic model assessment for insulin resistance
IL-6: interleukin-6
LDL: low-density lipoprotein
ML: machine learning
MUFA: monounsaturated fatty acids
NMR: nuclear magnetic resonance spectroscopy
P: particles
PL: phospholipids
PUFA: polyunsaturated fatty acids
SFA: saturated fatty acids
TG: triacylglycerols
VLDL: very low-density lipoprotein
XXL, XL, L, M, S: extra extra large, extra large, large, medium, small

## Data availability

Data described in the article, code book, and analytic code are held with the Department of Twin Research at King’s College London and will be made available using our normal procedures overseen by the Wellcome Trust and its guidelines as part of our core funding. The application is at: https://twinsuk.ac.uk/resources-for-researchers/access-our-data/

## References

1. Deelen J, Kettunen J, Fischer K, van der Spek A, Trompet S, Kastenmüller G, et al. A metabolic profile of all-cause mortality risk identified in an observational study of 44,168 individuals. Nat Commun 2019;10:3346.

2. Menni C, Fauman E, Erte I, Perry JR, Kastenmüller G, Shin SY, et al. Biomarkers for type 2 diabetes and impaired fasting glucose using a nontargeted metabolomics approach. Diabetes 2013;62:4270–6.

3. Menni C, Graham D, Kastenmüller G, Alharbi NH, Alsanosi SM, McBride M, et al. Metabolomic identification of a novel pathway of blood pressure regulation involving hexadecanedioate. Hypertension 2015;66:422–9.

4. Moayyeri A, Cheung CL, Tan KC, Morris JA, Cerani A, Mohney RP, et al. Metabolomic Pathways to Osteoporosis in Middle-Aged Women: A Genome-Metabolome-Wide Mendelian Randomization Study. J Bone Miner Res 2018;33:643–650.

5. Soininen P, Kangas AJ, Würtz P, Suna T, and Ala-Korpela M. Quantitative serum nuclear magnetic resonance metabolomics in cardiovascular epidemiology and genetics. Circ Cardiovasc Genet 2015;8: 192–206.

6. Ketema EB, Kibret KT. Correlation of fasting and postprandial plasma glucose with HbA1c in assessing glycemic control; systematic review and meta-analysis. Archives of Public Health 2015;73:1–9.

7. Sciarrillo CM, Koemel NA, Keirns BH, Banks NF, Rogers EM, Rosenkranz SK, et al. Who would benefit most from postprandial lipid screening? Clinical Nutrition 2021;40:4762–71.

8. Kolovou GD, Mikhailidis DP, Kovar J, Lairon D, Nordestgaard BG, Ooi TC et al. Assessment and clinical relevance of non-fasting and postprandial triglycerides: an expert panel statement. Cur Vasc Pharmacol 2011;9:258–70.

9. Blaak EE, Antoine JM, Benton D, Björck I, Bozzetto L, Brouns F, et al. Impact of postprandial glycaemia on health and prevention of disease. Obes Rev 2021;13:923–84.

10. Berry SE, Mills CE, Harding S, Bruce J, Gray R, Bapir M, et al. Lower postprandial lipemia after palmitic acid-rich fats with and without interesterification is associated with increased atherogenic lipoproteins versus a high MUFA oil (OR19-03-19). Curr Dev Nutr 2017;3:nzz046.OR19–03–19.

11. Berry SE, Valdes AM, Davies R, Al Khatib H, Delahanty L, Drew DA, et al. Large Inter-individual Variation in Postprandial Lipemia Following a Mixed Meal in over 1000 Twins and Singletons from the UK and US: The PREDICT I Study (OR19-06-19). Curr Dev Nutr 2017 9;3:nzz046.OR19-06-19.

12. Wildberg C, Masuch A, Budde K, Kastenmüller G, Artati A, Rathmann W, et al. Plasma metabolomics to identify and stratify patients with impaired glucose tolerance. J Clin Endocrinol Metab 2019;104:6357–6370.

13. Asnicar A, Berry SE, Valdes AM, Nguyen LH, Piccinno G, Drew DA et al. Microbiome connections with host metabolism and habitual diet from 1,098 deeply phenotyped individuals. Nat Med 2021;27:321–332.

14. Berry SE, Valdes AM, Drew DA, Asnicar F, Mazidi M, Wolf J, et al. Human postprandial responses to food and potential for precision nutrition. Nat Med 2020;26: 964–73.

15. Berry S, Drew DA, Linenberg I, Wolf J, Hadjigeorgiou G, Davies R, et al. Personalised REsponses to DIetary Composition Trial (PREDICT): an intervention study to determine inter-individual differences in postprandial response to foods. Protocol Exchange (2020).

16. Mazidi M, Valdes AM, Ordovas JM, Hall WL, Pujol JC, Wolf J, et al. Meal-induced inflammation: postprandial insights from the Personalised REsponses to DIetary Composition Trial (PREDICT) study in 1000 participants. The American journal of clinical nutrition 2021;114:1028–38.

17. Würtz P, Havulinna AS, Soininen P, Tynkkynen T, Prieto-Merino D, Tillin T, et al. Metabolite profiling and cardiovascular event risk: a prospective study of 3 population based cohorts. Circulation 2015;131:774–785.

18. Sampson, JN, Boca, SM, Shu, XO, Stolzenberg-Solomon, RZ, Matthews, CE, Hsing, AW et al. Metabolomics in epidemiology: sources of variability in metabolite measurements and implications. Cancer Epidemiol Biomarkers Prev 2013;22:631–640.

19. Floegel, A, Drogan, D, Wang-Sattler, R, Prehn, C, Illig, T, Adamski, J, et al. Reliability of serum metabolite concentrations over a 4-month period using a targeted metabolomic approach. PLoS One 2011;6:e21103.

20. Benjamini Y, Hochberg Y. Controlling the false discovery rate: a practical and powerful approach to multiple testing. R Stat Soc. 1995;57:289–300.

21. Xiao Q, Moore S. C, Boca S. M, Matthews, C. E, Rothman, N, Stolzenberg-Solomon, R. Z, et al. Sources of variability in metabolite measurements from urinary samples. PLoS One 2014;9:e95749.

22. Carayol M, Licaj I, Achaintre D, Sacerdote C, Vineis P, Key TJ, et al. Reliability of serum metabolites over a two-year period: a targeted metabolomic approach in fasting and non-fasting samples from EPIC. PloS one 2015;10:e0135437.

23. Secor, S. M. Specific dynamic action: a review of the postprandial metabolic response. Journal of Comparative Physiology B 2009;179:1–56.

24. Wallace M, Hashim YY, Wingfield M, Culliton M, McAuliffe F, Gibney MJ, et al. Effects of menstrual cycle phase on metabolomic profiles in premenopausal women. Human Reproduction 2010;25:949–56.

25. Nicholson, G, Rantalainen, M, Maher, A. D, Li, J. V, Malmodin, D, Ahmadi, K. R, et al. Human metabolic profiles are stably controlled by genetic and environmental variation. Mol. Syst. Biol 2011;7:525.

26. Wojczynski MK, Glasser SP, Oberman A, Kabagambe EK, Hopkins PN, Tsai MY, et al. High-fat meal effect on LDL, HDL, and VLDL particle size and number in the Genetics of Lipid-Lowering drugs and diet network (GOLDN): an interventional study. Lipids Health Dis 2011;10:181.

27. Sarwar N, Danesh J, Eiriksdottir G, Sigurdsson G, Wareham N, Bingham S, et al. Triglycerides and the risk of coronary heart disease: 10,158 incident cases among 262,525 participants in 29 Western prospective studies. Circulation 2007;115:450–8.

28. Johansen MØ, Nielsen SF, Afzal S, Vedel-Krogh S, Smith GD, Nordestgaard BG. Very low-density lipoprotein cholesterol may mediate a substantial component of the effect of obesity on myocardial infarction risk: The Copenhagen General Population Study. Clin Chem 2021;67: 276–287.

29. Karpe F, Hellénius ML, Hamsten A. Differences in postprandial concentrations of very-low-density lipoprotein and chylomicron remnants between normotriglyceridemic and hypertriglyceridemic men with and without coronary heart disease. Metabolism 1999;48:301–7.

30. Mills CE, Harding SV, Bapir M, Mandalari G, Salt LJ, Gray R, et al. Palmitic acid–rich oils with and without interesterification lower postprandial lipemia and increase atherogenic lipoproteins compared with a MUFA-rich oil: A randomized controlled trial. Am J Clin Nutr, 2021;113:1221–1231.

